# Kaptive Web: user-friendly capsule and lipopolysaccharide serotype prediction for *Klebsiella* genomes

**DOI:** 10.1101/260125

**Authors:** Ryan R Wick, Eva Heinz, Kathryn E Holt, Kelly L Wyres

**Affiliations:** Department of Biochemistry and Molecular Biology, Bio21 Molecular Science and Biotechnology Institute, University of Melbourne, Victoria, Australia; Wellcome Trust Sanger Institute, Hinxton, UK

## Abstract

As whole genome sequencing becomes an established component of the microbiologist’s toolbox, it is imperative that researchers, clinical microbiologists and public health professionals have access to genomic analysis tools for rapid extraction of epidemiologically and clinically relevant information. For the gram-negative hospital pathogens such as *Klebsiella pneumoniae*, initial efforts have focused on detection and surveillance of antimicrobial resistance genes and clones. However, with the resurgence of interest in alternative infection control strategies targeting *Klebsiella* surface polysaccharides, the ability to extract information about these antigens is increasingly important.

Here we present Kaptive Web, an online tool for rapid typing of *Klebsiella* K and O loci, which encode the polysaccharide capsule and lipopolysaccharide O antigen, respectively. Kaptive Web enables users to upload and analyse genome assemblies in a web browser. Results can be downloaded in tabular format or explored in detail via the graphical interface, making it accessible for users at all levels of computational expertise.

We demonstrate Kaptive Web’s utility by analysis of >500 *K. pneumoniae* genomes. We identify extensive K and O locus diversity among 201 genomes belonging to the carbapenemase-associated clonal group 258 (25 K and six O loci). Characterisation of a further 309 genomes indicates that such diversity is common among the multi-drug resistant clones and that these loci represent useful epidemiological markers for strain subtyping. These findings reinforce the need for rapid, reliable and accessible typing methods such as Kaptive Web.

Kaptive Web is available for use at kaptive.holtlab.net and source code is available at github.com/kelwyres/Kaptive-Web.

## Introduction

Whole genome sequencing (WGS) represents a powerful tool for characterisation and public health surveillance of bacterial pathogens. This technology is now routinely used by a number of public health laboratories (1–3), and there is increasing interest in its use in clinical labs (4–6). While there are several well developed protocols which use WGS data for determination of multi-locus sequence types (STs), resistance gene profiling and phylogenetic investigations (7–9), there remain gaps in the repertoire, e.g. characterisation of species-specific antigens is currently restricted to a small number of species (10–12). Furthermore, many WGS analyses rely on software via a command line interface that require bioinformatics skills to install, execute and interpret, thereby limiting their accessibility. Instead, we need tools that can extract information and present it in an easily interpretable manner to bioinformaticians, public health professionals and clinicians alike (5, 7, 8).

*Klebsiella pneumoniae* is a major cause of healthcare-associated infections with high rates of multi-drug resistance (MDR) (13). In particular, the emergence and global dissemination of extended-spectrum beta-lactamase (ESBL) and carbapenemase producing (CP) clones is a major concern and has led to the recognition of *K. pneumoniae* as an urgent public health threat (14, 15). With the lack of new antimicrobial therapies, there has been a resurgence of interest in alternative strategies such as phage therapy (16–19), monoclonal antibody therapy (20–23) and vaccination (24–26). Several therapeutic targets have been suggested, and the polysaccharide capsule (K antigen) and lipopolysaccharide (O antigen) are among the most frequent. Both are also considered key virulence determinants that are necessary to establish infection, primarily owing to their serum resistance and anti-phagocytic properties (27–31). Of note, capsular serotypes vary substantially in the degree of serum resistance they provide. For example, K1, K2 and K5 are highly serum resistant and are associated with hypervirulent strains that differ from classical *K. pneumoniae* in that they commonly cause community-acquired disease (32–34). Despite the importance of these loci, *K. pneumoniae* serotyping is not widely available, even in large central public health laboratories, and the most practical option for most laboratories is genotyping the loci involved in antigen biosynthesis via multiplex PCR (35, 36) or WGS (37, 38).

The lipopolysaccharide comprises three subunits: lipid A, the core oligosaccharide and the O antigenic polysaccharide (39). With only one exception, the key determinants of the O antigenic polysaccharide are co-located at the O locus (previously known as the *rfb* locus) (36, 40–42). While 10 serologically distinct O antigens have been recognised, many isolates are non-typeable (23, 43) and investigations have identified 12 distinct O loci (25, 36). Interestingly, both the O1 and O2 antigens, which are by far the most common (23, 25, 43), are each associated with the same two loci, O1/O2v1 and O1/O2v2 (25). Expression of the v1 locus results in production of D-galactan I, characteristic to a subset of O2 antigens (41, 44). Expression of the v2 locus results in production of D-galactan III associated with the remaining O2 subtypes (21, 45). Regardless of subtype, any O2 antigen can be converted to O1 by the addition of D-galactan II, which requires the products of *wbbY* and *wbbZ* that are located outside of the O locus, i.e. elsewhere in the genome (44).

The *Klebsiella* polysaccharide capsule is produced through a Wzy-dependent process (46), for which the synthesis and export machinery are encoded in a single 10-30 kbp region of the genome known as the K locus (47, 48). Seventy-seven distinct capsule phenotypes have been recognised by serological typing (49), but many isolates are serologically non-typeable. We recently explored the K loci among a large diverse *K. pneumoniae* WGS collection and were able to define 134 distinct loci on the basis of protein coding gene content, suggesting there are at least this many distinct capsule types circulating in the population (38).

Given the interest in targeting these diverse surface polysaccharides, and the lack of accessible serotyping assays for *K. pneumoniae*, tools for WGS-based K and O locus typing will be essential for researchers, clinicians and public health microbiologists. We previously developed Kaptive for K locus typing from WGS assemblies (38), which has become a key component of the *Klebsiella* bioinformatics tool kit (50–52), but it requires command-line skills and a degree of bioinformatics expertise to use. Here we present Kaptive Web, an easy-to-use web-based implementation of the Kaptive algorithm which has been extended to type both K and O loci. We demonstrate its utility: (i) for the identification of K and O loci for serotype prediction and as epidemiological markers; and (ii) to inform the design and implementation of control strategies targeting the capsules or lipopolysaccharides of *K. pneumoniae.*

## Results and discussion

### Introducing Kaptive Web

Kaptive Web is a browser-based method for running Kaptive and visualising the results. Users upload one or more assemblies and select their preferred typing database, after which command-line Kaptive is automatically run on the remote server (**Fig 1**). Results appear in a table with one row per genome assembly showing key details: the best-matching locus from the reference database, the match confidence, nucleotide identity and coverage compared to the reference (**Fig S1**). Rows are coloured based on match confidence, with six possible levels:

**Figure 1:**
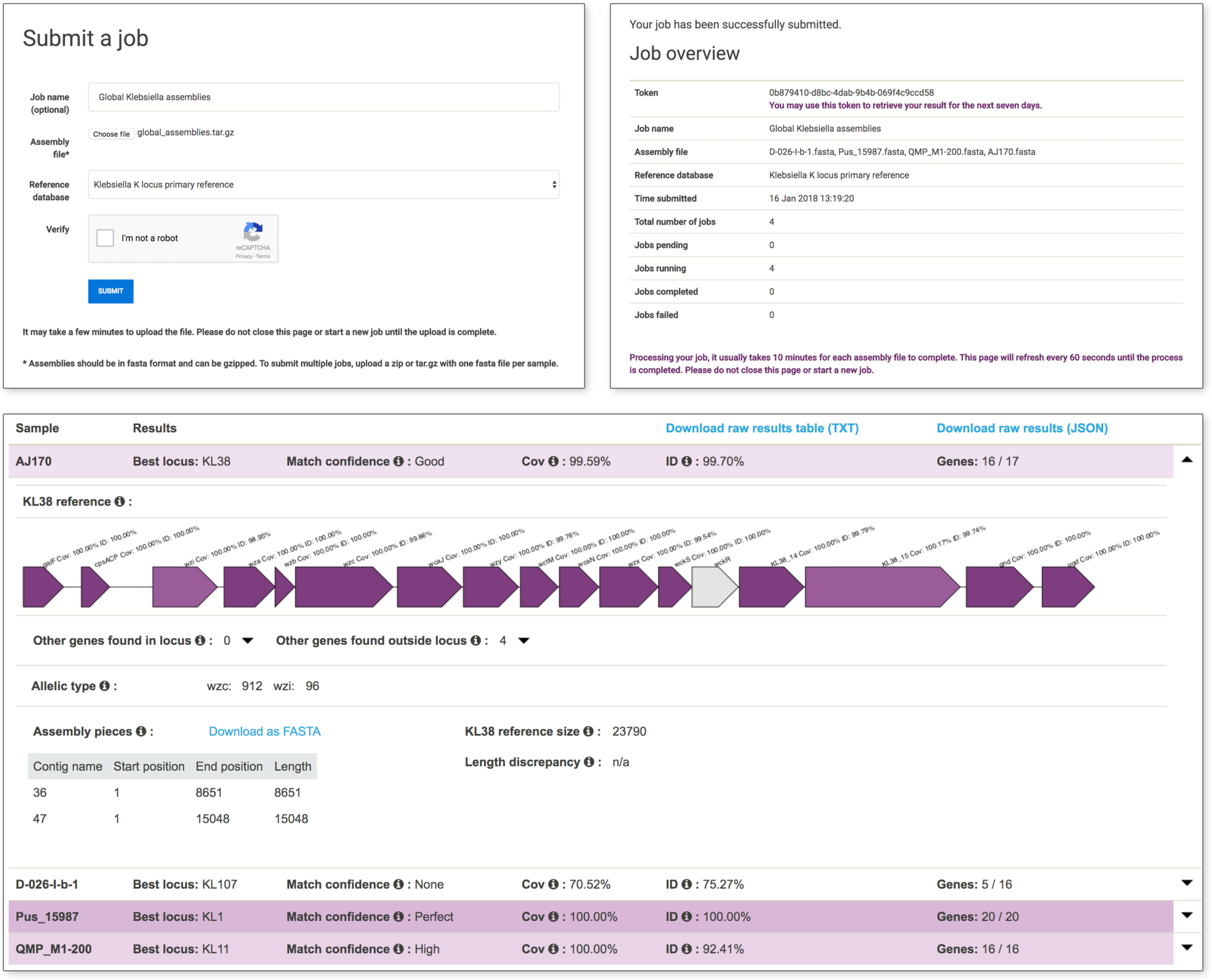
Kaptive Web screenshots. Top left: users choose a file or zipped directory on their local computer and select the desired reference database. Top right: job information screen detailing the number of assemblies uploaded and their status. Bottom: Kaptive Web results for four *Klebsiella* isolates, with the first result expanded.

Perfect | The locus was found in a single piece with 100% coverage and 100% nucleotide identity to the reference.
Very high | The locus was found in a single piece with ≥99% coverage and ≥95% nucleotide identity to the reference, with no truncated/missing genes and no extra genes compared to the reference.
High | The locus was found in a single piece with ≥99% coverage, with ≤3 truncated/missing genes and no extra genes compared to the reference.
Good | The locus was found in a single piece, or with ≥95% coverage, with ≤3 truncated/missing genes and ≤1 extra gene compared to the reference.
Low | The locus was found in a single piece, or with ≥90% coverage, with ≤3 truncated/missing genes and ≤2 extra genes compared to the reference.
None | Did not qualify for any of the above

Clicking on an assembly row expands the view to show more detail, including a diagram of the best-matching locus with genes coloured by tBLASTn coverage and identity (**Fig 1**). Beneath the locus diagram are two expandable lists of additional K or O locus genes identified inside and outside the locus region of the query genome (genes that are not usually present in the reference locus). It is common to see matches to a small number of additional genes outside the locus region of the query due to sequence homology (see (38) for further details). An additional gene within the locus region of the query genome may indicate that the genome has a novel locus type and will likely correspond to a large length discrepancy from the reference (shown on the right side of the display). In such a case, users may wish to perform further analyses outside of Kaptive Web. To facilitate this, Kaptive Web lists the position of the locus in the query genome (shown on the left side of the display), along with a link that allows these assembly regions to be downloaded in FASTA format. For K locus typing, Kaptive Web will also report the alleles for the conserved *wzc* and *wzi* K locus genes, to allow compatibility with earlier schemes that focused on these genes (37, 53, 54).

**Fig 1** shows Kaptive Web K locus results for four isolates from a global *Klebsiella* diversity study dataset (55), with varying degrees of data quality. The assembly for strain Pus_15987 has a ‘Perfect’ match for KL1. Strain D-026-I-b-1 has a best matching locus of KL107, though poor assembly quality has resulted in very low identity and coverage, and consequently a confidence level of ‘None’. Strain QMP_M1-200 has a ‘High’ match for KL11. It contains the entire KL11 sequence, but with a moderate amount of divergence (92% nucleotide sequence identity). In most cases, minor nucleotide divergence likely does not affect capsule phenotype. However, it should be noted that even a single nonsense or frame-shift mutation can have important implications; e.g. the K22 and K37 capsules are encoded by loci distinguished only by a nonsense mutation in the acetyltransferase gene (48). Kaptive Web clearly identifies potential nonsense or frame-shift mutations by marking such loci with ‘missing genes’.

The results for strain AJ170 are expanded in **Fig 1**, showing the full Kaptive Web visualisation. It has a very good coverage and identity match to KL38, yet the locus was not found in a single contiguous piece of the assembly. Kaptive was also unable to find a translated protein sequence for one of the KL38 genes, *wckR*, illustrated by the grey colouring in the locus diagram. In this instance, both issues (discontiguous locus sequence and missing gene) were caused by a break in the assembly, splitting the locus over two contigs. This may have resulted from poor read coverage, in which case *wckR* may have been intact and functional in the original isolate.

Alternatively, the assembly break may have resulted from an insertion sequence interrupting *wckR*, in which case gene function is likely lost; such interruptions have been characterised in a number of *Klebsiella* K loci (38). This uncertainty is why AJ170 only achieved a ‘Good’ confidence score for KL38.

### O-locus database and typing

The Kaptive algorithm was originally developed and validated for typing the K locus of *K. pneumoniae* (38), but it can in principle be used to type any variable locus that occurs no more than once per genome. In Kaptive Web, we apply it to O locus typing, which follows mostly the same procedure as K locus typing but with two unique aspects. First, serological types 01 and 02 are distinguished not by the 0 locus but by two genes, *wbbY*and *wbbZ*, elsewhere in the chromosome. Second, the 0 locus shared by the 01 and 02 serotypes comes in two varieties (v1 and v2), which are distinguishable using genomic data but are serologically cross-reactive.

These aspects are incorporated into Kaptive as follows: (i) the relevant 0 locus variant is reported as v1 or v2; (ii) an additional search is conducted for *wbbY* and *wbbZ* to decide whether the locus should be reported as 01 or 02. If only one of *wbbY* or *wbbZ* is found, Kaptive will give a label of 01/02 (i.e. possibly either).

A recent study of 03 antigens identified several subtypes that can be distinguished serologically and genotypically (03, 03a and 03b, (20)). The 03 and 03b loci correspond to the 03l and 03s loci previously described in Follador *et al.* (25) and are distinguished by divergent sequence of the *wbdA* and *wbdD* genes (25). The 03 and 03a loci are distinguished by a single point mutation in *wbdA* (C80R) (20). Kaptive is able to distinguish the former pair but not the latter, and therefore designates 03 loci as either 03/03a or 03b.

We assessed the accuracy of Kaptive 0 locus typing by application to the *Klebsiella* global diversity study genomes (55) for which 0 locus types were previously inferred on the basis of nucleotide variation in the universally conserved *wzm* and *wzt* genes (25). 0f the 309 WGS assemblies, the number which matched each confidence level were: 3 ‘Perfect’, 212 ‘Very high’, 28 ‘High’, 56 ‘Good’, 3 ‘Low’ and 7 ‘None’ (**Table S1**). The assemblies with a confidence of ‘Low’ or ‘None’ were possibly due to low assembly quality; seven had the 0 locus split over multiple contigs and three had very low coverage. There was very good agreement between the 0 locus types defined previously (25) and the Kaptive results - only 8/309 assemblies had discrepancies. 0f those, four were cases where the previous type was 01 and Kaptive assigned 01/02 (i.e. it only found one of *wbbY* and *wbbZ)*, and one where the previous type was 01 and Kaptive assigned 02. The remaining three discrepancies were all between 03 and 0L104 which are distinguished by their *wbdD* genes (25). Isolates AJ031 and D-026-I-b-1 were mistyped due to poor assembly - not all of the 0 locus was represented. Isolate U_13792_2 was typed as 03s by Kaptive but previously assigned 0L104, and manual inspection of the predicted WbdD amino acid sequence suggested that this strain produces a hybrid WbdD protein. Given the more subtle distinctions between the 03/03a, 03b and 0L104 loci, we further tested the accuracy of Kaptive’s 0 locus typing using 13 additional 03 *K. pneumoniae* for which genome data and 0 antigen phenotypes were previously determined (20), and found Kaptive correctly typed all 13 genomes (**Table S2**).

### Application of Kaptive Web to track K and O locus diversity in MDR clones

Increasing rates of antimicrobial resistance, particularly against last-line drugs such as carbapenems, has led to a resurgence of interest in phage and monoclonal antibody therapies and vaccinations targeting *K. pneumoniae* (16–25). There is particular interest in targeting the globally distributed MDR clones including CG258 (21-23, 26). ST258, the most well-known member of CG258, has rapidly become the most common cause of CP *Klebsiella* infections in the United States (56). Recent studies suggest low lipopolysaccharide diversity in this clone, with the majority of ST258 isolates expressing the 02 antigen (21, 23). Similarly, early studies reported that ST258 harboured just two distinct capsule types (57). However, subsequent work has shown greater K locus variation (58), particularly among other members of the clonal group, e.g. ST11, ST340 and ST437, that are frequent causes of CP infections outside of the United States (59). To explore the broader diversity of K and O loci in CG258, we downloaded all CG258 genome assemblies available in GenBank (n=201 as at 12 Oct 2017, see **Table S3**) and analysed them using Kaptive Web. ‘Good’ or better K and O locus calls were obtained for 173 (86%) and 186 (93%), respectively (**Table S3**). Importantly, while there were dominant types (KL107 and O2v2, the combination of which accounted for 65 (32%) of isolate genomes), there was also much diversity: 25 K loci and six O loci in total (not distinguishing O1 and O2, see **Fig 2**).

**Figure 2:**
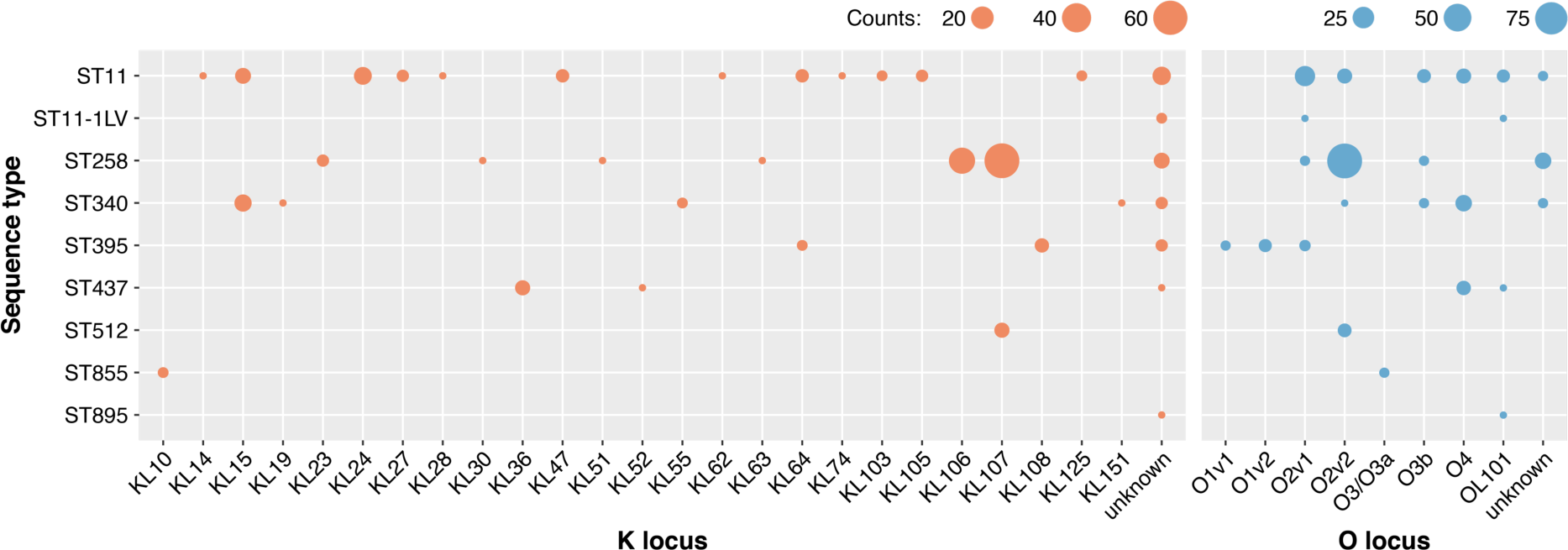
K and O locus diversity among 201 CG258 genomes. Counts of genomes representing each K locus (orange) and O locus (blue) are shown by multi-locus sequence type (ST). ST11-1LV indicates an unassigned single locus variant of ST11. Genomes with ‘Low’ or ‘None’ confidence Kaptive calls are shown as ‘unknown.’

There is now emerging evidence that like CG258, other globally distributed MDR clones also harbour diverse K and O loci (50, 60). For example, **Fig 3** shows the core chromosomal phylogeny of the 309 global genomes (55), highlighting Kaptive Web’s K and O locus calls for three additional globally distributed MDR lineages (55, 59). Locus exchange tends to result from recombination and demarcates diverging sublineages within the expanding MDR clones (57, 58), hence the locus calls can also serve as epidemiological markers for subtyping of MDR strains (54, 57). Therapies specific to the dominant K or O antigens may apply further selective pressure that shifts the population towards different types, as is well documented following the introduction of protein-conjugate vaccines targeting the *Streptococcus pneumoniae* polysaccharide capsule (61, 62). The success of new control measures directed at K and O loci will therefore depend on reliable tracking of K and O loci in the *K. pneumoniae* population. Kaptive and Kaptive Web provide simple WGS-based solutions to monitor these trends and ensure that therapies are well-targeted and keep up with *K. pneumoniae* evolution. With its graphical interface and remote computation, Kaptive Web also makes these analyses accessible to the wider public health community.

**Figure 3:**
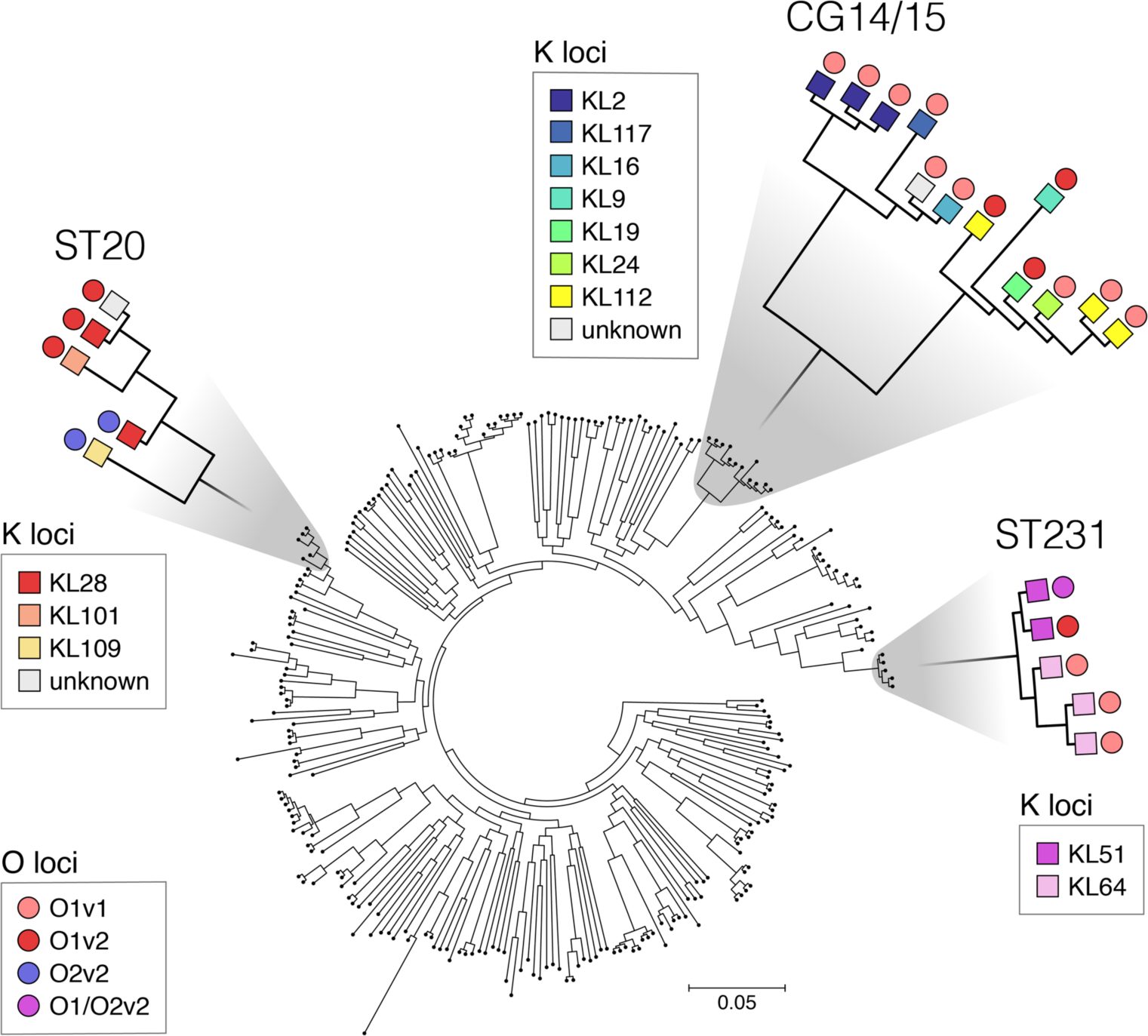
Phylogenetic tree of global *K. pneumoniae* isolates with Kaptive results for K and O loci noted on selected clonal groups. Only Kaptive locus calls with a confidence of ‘Good’ or better were included in this figure. Lower confidence matches were labelled ‘unknown’.

## Materials and Methods

### O locus definitions

Unlike K loci which were defined on the basis of gene content, O loci have been defined by sequence identity in the conserved *wzm* and *wzt* genes (25). For example, two O loci can have the same genes, but modestly divergent sequence (>5% divergence). Kaptive is compatible with these definitions because it first chooses a best locus based on a nucleotide search. Only then does it tally the gene content of the locus.

A complication comes from the fact that O antigens O1 and O2 are encoded by the same two O loci. It is the presence or absence of two other genes elsewhere in the genome, *wbbY* and *wbbZ*, which determines the specific antigen. When both genes are present, D-galactan II is produced leading to the O1 antigen. When they are absent, the O2 antigen results. We have added the appropriate logic to Kaptive (both command-line Kaptive and Kaptive Web) so it will report the locus as O1 or O2 based on the presence/absence of these genes as determined by tBLASTn search with coverage and identify thresholds of 90% and 80%, respectively. If Kaptive finds only one of the two genes, it will report the locus as O1/O2.

A Kaptive-compatible O locus reference database comprising the O loci reported in (25) plus the O8 locus (Genbank accession: AB819963.1) is available for download via the command-line Kaptive GitHub page (github.com/katholt/Kaptive).

### Kaptive Web

Kaptive Web is available for use at kaptive.holtlab.net. It was developed using the web2py framework (63). The source code for the web implementation is available on GitHub, so users can host their own copy of the software (github.com/kelwyres/Kaptive-Web). Kaptive Web automatically populates the ‘Reference database’ selection with the contents of command-line Kaptive’s database directory, allowing automatic compatibility with any new locus databases for *Klebsiella* or other bacteria.

### Genome data for K and O locus characterisation

Sequence read data for 309 *K. pneumoniae* sequenced as part of the global diversity study (55) and 13 03 antigen-producing isolates (20) were assembled *de novo* using Unicycler v0.4.1 (64). Genome assemblies were uploaded to Kaptive Web in a single compressed data directory and analysed with the *Klebsiella* primary K locus and the *Klebsiella* 0 locus databases. Kaptive Web utilises a 16 core 64 GB RAM server hosted by the Australian NeCTAR cloud. A single genome analysis requires approximately 4 minutes and 20 seconds to complete for the K and 0 locus databases, respectively. Kaptive Web runs multiple genomes in parallel, so the total analysis times for the global dataset were 52 minutes (K locus) and 12 minutes (0 locus). The results were inspected via the Kaptive Web graphical interface and downloaded in tabular format (see **Table S1**).

The same protocol was used for characterisation of 201 publicly available CG258 genome assemblies (see **Table S3**). These genomes were identified among the complete set of *Klebsiella* genomes (downloaded from GenBank on 12 0ct 2017) on the basis of ST information generated using Kleborate (github.com/katholt/Kleborate). STs 11, 258, 340, 395, 437, 512, 855, and 895 were included in the analyses.

## Acknowledgements

Web development services were provided by the eResearch group at the University of Melbourne.

